# SARS-CoV-2 spike protein amyloid fibrils impair fibrin formation and fibrinolysis

**DOI:** 10.1101/2025.06.30.661938

**Authors:** Henrik Westman, Per Hammarström, Sofie Nyström

## Abstract

Long COVID, also known as post-acute sequelae of COVID-19 (PASC) from SARS-CoV-2 infection, is a debilitating and persistent disease of multiple systems and organs. WHO reports a total of 778 million registered SARS-CoV-2 infections as of June 2025. Recent studies indicate that as many as 25% of COVID patients will experience at least one symptom of Long COVID. Long COVID pathophysiology is a complex and not fully established process. One prevailing theory is that the formation of fibrin amyloid microclots (fibrinaloids), due to SARS-CoV-2 infection, can induce persistent inflammation and capillary blockage. An association between the amyloidogenic Spike protein of SARS-CoV-2 and impaired fibrinolysis has previously been made when it was observed that fibrin clots formed in the presence of a mixture of amyloid fibrils from the spike protein displayed a resistance to plasmin-mediated lysis. Here we investigated the molecular processes of impaired fibrinolysis using seven amyloidogenic SARS-COV-2 Spike peptides. Five out of seven Spike amyloid fibrils appeared not to substantially interfere with the fibrinogen-fibrin-fibrinolysis process in vitro, while two spike fibrils were active in different ways. Spike601 amyloid fibrils (sequence 601-620) impaired thrombin mediated fibrin formation by binding and sequestering fibrinogen but did not affect fibrinolysis. On contrary fibrin clots formed in the presence of Spike685 amyloid fibrils (sequence 685-701) exhibited a marked resistance to plasmin mediated fibrinolysis. We conclude that Spike685 amyloid fibrils can induce dense fibrin clot networks, as well as incorporate fibrin into aggregated structures that resist fibrinolysis. These results demonstrate how the Spike protein of SARS-CoV-2 could contribute to the formation fibrinolysis-resistant microclots observed in long COVID.

## Introduction

During the Coronavirus disease 2019 (COVID-19) pandemic caused by the Severe acute respiratory syndrome coronavirus 2 (SARS-CoV-2), it was reported that many patients, recent studies say up to 25%, exhibited physical symptoms several months after cleared virus infection [1] [2] [3] [4]. These symptoms have been observed to manifest or persist as a wide range of non-specific clinical symptoms involving the entire body, affecting multiple organs and systems [5] [6] [7]. Furthermore, patients commonly exhibit chronic fatigue and cognitive dysfunction, commonly known as “brain fog”, along with a majority showing post-exertional symptom exacerbation (PESE). This has led to the classification of a disease termed “long COVID”, “post-COVID conditions”, or “post-acute sequelae of COVID-19 (PASC)”.

Regarding these multi-system effects of long COVID, one discovery that could explain the collection of varied symptoms are formation of microclots in the blood of patients [5] [8]. These microscopic blood clots (or thrombi) have been shown to consist of misfolded, amyloid forms of the clotting protein fibrin, as well as an entrapment of several other proteins. Because of their size and composition, microclots have been suggested to block microcapillaries, limit O_2_ transfer to tissues, induce oxidative stress, and promote release of inflammatory cytokines. Another aspect of microclots is the resistance to proteolytic breakdown known as fibrinolysis. Therefore, capillary blockages by microclots that remain in the body over time and has been argued to contribute and play a role in the pathophysiology of long COVID.

More specifically it has been proposed that the Spike protein of SARS-CoV-2 virus cause the clotting pathology of long COVID. The Spike protein can both activate clotting factors [9] and have amyloidogenic properties [10]. By interacting with the fibrinogen protein, inducing inflammation, and affecting coagulation and fibrinolytic processes, the SARS-CoV-2 Spike protein could give rise to an “amyloid-mediated impaired fibrinolysis” [9] [11]. The association of misfolded, amyloidogenic proteins to altered blood coagulation and fibrinolysis has also been recognised in other amyloid diseases, such as Cerebral amyloid angiopathy (CAA) of amyloid beta (Aβ) [12] [13]. The mechanism of which has recently been proposed to be mediated by Aβ1-42 protofibrils making complexes with fibrinogen [14] [15]

SARS-CoV-2 is an enveloped positive-sense single-strand RNA virus with the virion particle composed of 4 structural proteins [16] [17]. These proteins are the Spike, envelope, membrane, and nucleocapsid proteins. Each monomer of the large homotrimeric Spike protein consists of two subunits (S1 and S2), where S1 contains a domain with the receptor binding motif that enables viral entry [16]. This domain of the SARS-CoV-2 Spike protein exhibits the highest amount of mutational rate. The Spike protein has been characterized as being an amyloidogenic protein [10]. Upon *in vitro* cleavage of the full-length Spike protein by neutrophil elastase and subsequent incubation, networks of branched amyloid fibrils were produced. We previously identified 7 amyloidogenic peptide sequences within the Spike protein. Amyloid fibrils composed of a mixture of these seven peptides impaired the fibrinolysis process *in vitro* [10].

The interest from the scientific community for the hypothesis that formation of persistent amyloid-like small thrombi or microclots is mechanistically associated with amyloidogenisis of SARS-CoV-2 Spike protein [18] rendered us to investigate this link in detail. We specifically asked if amyloid fibrils from different Spike sequences influenced the fibrinogen-fibrin-fibrinolysis pathway using pure biochemical systems.

## Results

We first revisited our protocol for formation of amyloid fibrils from synthetic peptides of the SARS-CoV-2 Spike protein (original Wuhan sequence) (**Fig. 1**) [10]. At a peptide concentration of 0.5 and 1 mg/ml all seven peptides formed amyloid fibrils as deduced by ThT fluorescence intensity and negative stain transmission electron microscopy (TEM) (**Fig. 1**) as well as Thioflavin T (ThT) kinetics assay and Congo red binding and birefringence (**SFig. 1**).

**Figure 1.**
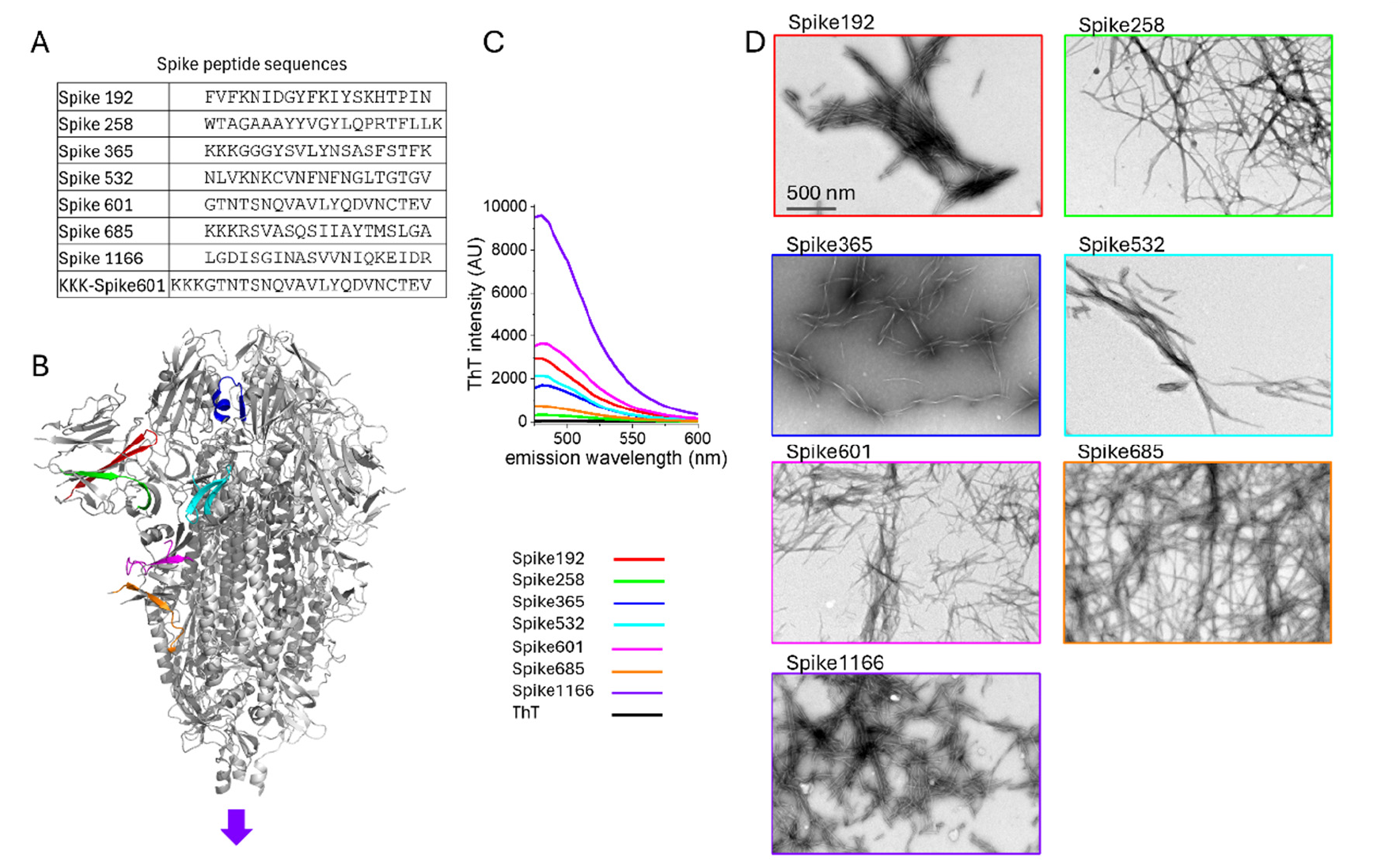
Amyloidogenic sequences in SARS-CoV-2 Spike and corresponding peptides (A, B) were selected by WALTZ prediction [19]and their amyloid properties were tested by ThT intensity measurements (C) and TEM analysis (D). Fibrils were formed at 0.5 mg/ml of Spike peptide within 24 h. Additional verification methods for amyloid are found in SFig 1. The structure model (in B) displays the location of the amyloidogenic segments within one protomer of the prefusion state of the Spike protein trimer determined by Cryo-EM (PBD: 6VXX)[20].

### Fibrin formation and fibrinolysis

A procedure for thrombin-induced fibrin clot formation and plasminogen/tPA fibrinolysis was established. Addition of thrombin to fibrinogen in a well of a 96-well plate instantly induced fibrin formation leading to the formation of light scattering fibrin fibers (**Fig. 2A**). At 60 minutes content of the well has formed a polymerized gel or clot. The formed clot was thereafter covered with a solution of plasminogen and catalytic amounts of tissue plasmin activator (tPA), generating plasmin and the clot is slowly dissolved. The breakdown of the fibrin clot is monitored as loss of light scattering until baseline was reached (**Fig. 2A**).

**Figure 2.**
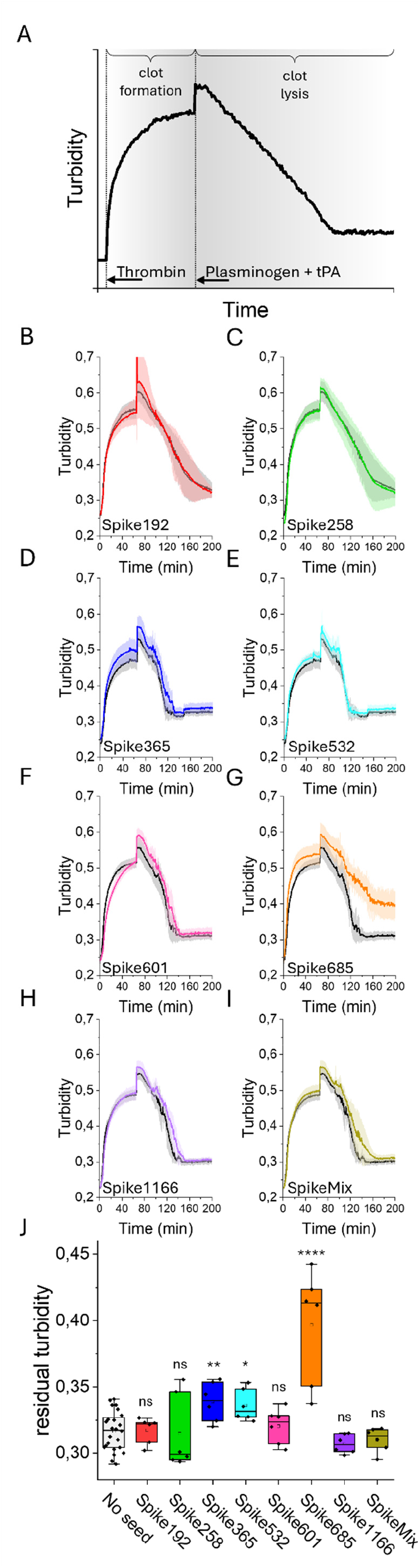
A) Turbidity curve of thrombin-induced fibrin polymerization. Clot formation was performed at a concentration of 0.5 mg/ml fibrinogen and 0.2 U/ml thrombin, a final volume of 75 µl. When the reaction reached its plateau, fibrinolysis was induced by addition of 75 µl 1.0 µM plasminogen and 2,4 nM tPA. Reactions were performed at 37 °C and turbidity was measured as OD at 355 nm. B-I) Clot formation and lysis were monitored in the absence and presence of Spike amyloid fibrils (final concentration 10 µg/ml) with one unseeded and two seeded experiments in each plate. Each experiment was run in 6 replicates. The graphs show the average of each spike amyloid fibril seeded experiment in color with its corresponding control, from the same plate, in grey. The shaded area depicts standard deviation. J) Residual turbidity at 200 minutes for all control experiments (no seed) (n=24) and each spike seeded sample (n=6) Two-sample T-test was performed: p>0,05=*; p>0,005=**; p>0,0001=****.

Based on experimental limitation of the fast initial rate of clot formation (first 5-10 minutes) the assay was performed using replicates of two Spike peptide amyloids where 10 µg/ml of Spike amyloid fibrils was added to each well, containing 625 µg/ml fibrinogen. One control sample, containing no addition of amyloid was run side-by-side in each plate (**Fig. 2 B-E**). Clot formation was monitored for 60 minutes. We found that Spike601 amyloid fibrils was the only Spike amyloid with a tendency to delay fibrin clot formation while addition of Spike365 and Spike685 amyloid fibrils resulted in higher turbidity at the plateau (**Fig. 2D,G**). Clot lysis was then initiated by addition of plasminogen and tPA. Spike685 amyloid fibril addition resulted in extensive residual turbidity at the experimental endpoint (**Fig. 2G**). Spike mix amyloids formed when co-aggregating the seven Spike peptides, also rendered a slight delay of fibrin clot lysis (**Fig 2E**) concomitant with our previous results [10] but no residual turbidity at endpoint 200 min was detected. A quantitative comparison of lysis efficacy between all Spike seeded reactions relative to a pool of all control experiments showed that presence of Spike685 amyloid fibrils during fibrin formation significantly impaired lysis of the formed clot (**Fig 2J**). This was also observed to some extent for Spike365 and Spike532 amyloid fibrils (**Fig 2J**), but not for Spike 192 amyloid fibrils at this concentration of seed.

The diverging behavior of Spike601 and Spike685 amyloid fibrils on the fibrinogen-fibrin-fibrinolysis process urged us to make concentration dependent experiments of these seeds. A titration of Spike601 and Spike685 amyloid fibrils in the turbidity assay was hence performed. Delayed fibrin formation in presence of Spike601 amyloid fibrils and, orthogonally, an increase in turbidity upon addition of Spike685 amyloid fibrils was detected (**Fig 3A,B,C**). For the fibrinolysis addition of Spike685 amyloid fibrils resulted in a highly significant concentration dependence on residual turbidity, showing that fibrinolysis became increasingly incomplete as the concentration of the Spike685 amyloid fibrils was increased (**Fig 3B,D**). For Spike601 amyloid fibrils on the other hand, only miniscule increase in residual turbidity was observed with higher concentration of amyloid (**Fig 3A,D**) showing that this is a specific effect for Spike685 amyloid fibrils.

**Figure 3.**
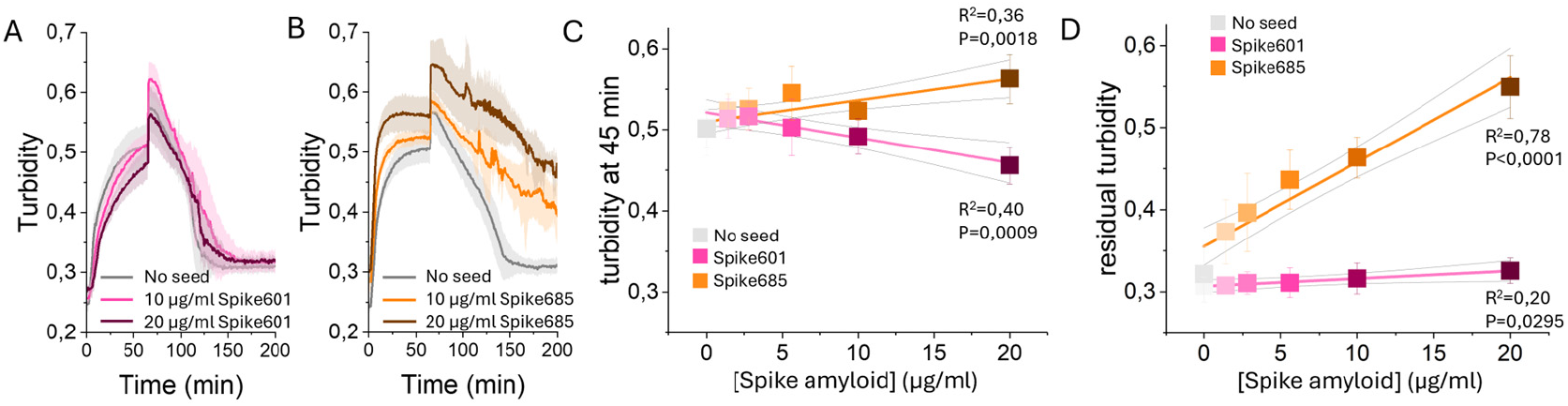
A, B) Concentration dependent impact of Spike601 and Spike685 amyloid fibrils respectively present during fibrin formation, exemplified by samples run with 0, 10 and 20 µg/ml Spike amyloid addition. C) Quantitative analysis of clot formation time at 45 minutes. D) Quantitative analysis of clot lysis measured as residual turbidity at 200 minutes. Average of four replicates with standard deviation. Colored lines in C and D represent linear regression fits to test trend significance (p values <0.05) with associated R square. The grey lines show 95% confidence intervals for the fit.

Spike 365 and Spike 685 amyloid fibrils were initially produced with a triple-lysine (KKK) solubility tag [10] (**Fig 1A**). This feature was exploited by using the tag as an amine rich tag for covalent NHS coupling of the fluorescent marker Cy5 on preformed fibrils. Subsequently we also made Spike601 peptide with a KKK-tag and labeled fibrils with Cy5. In combination with fluorescein labeled fibrinogen, we could quantify the co-precipitation of these three Spike amyloid fibrils and fibrinogen before thrombin cleavage and the residual fibrin in presence of the respective Spike amyloid fibrils after plasmin lysis of thrombin induced clots by spectroscopy using hyperspectral microscopy [21]. Spike365 amyloid fibrils did not pull-down fibrinogen or hamper fibrinolysis (**Figure 4A**). Fibrinogen co-precipitated with Spike 601 amyloid fibrils to a significantly higher degree than it did with either of Spike 365 and Spike685 amyloid fibrils and the co-localization of fluorescence intensities were retained also after lysis (**Figure 4B, D**). Co-precipitation of fibrinogen with Spike685 amyloid fibrils was similar to that of Spike365 amyloid fibrils (**Fig 4A, C, D**). Most strikingly, presence of Spike685 amyloid fibrils during fibrin clot formation resulted in a fibrinolysis product where residual fibrin and Spike amyloid co-localized at the resolution of this technique which is 25 µm (**Figure 4 C, D**).

**Figure 4.**
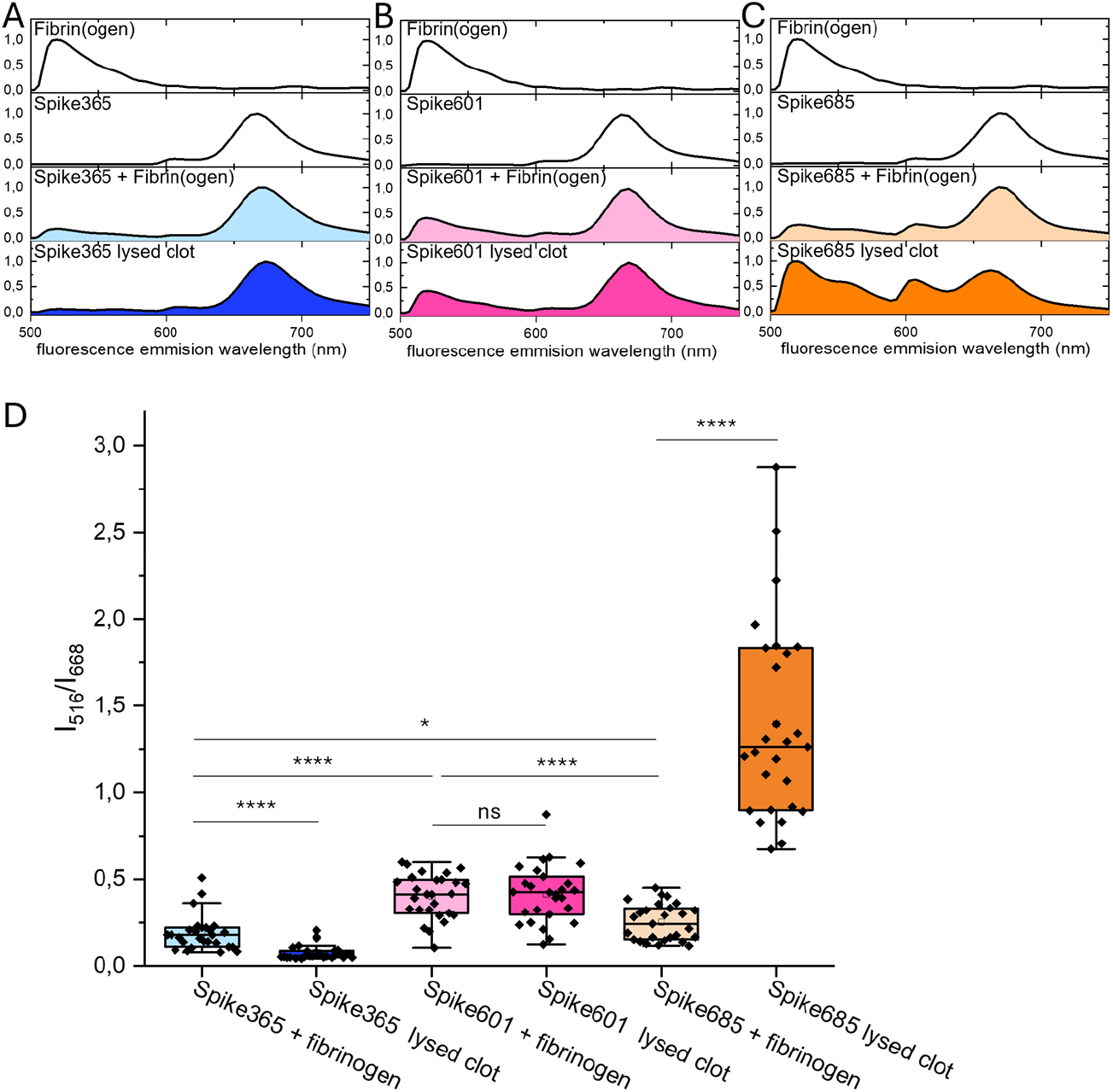
Fluorescence hyperspectral microscopy was performed for spectroscopic analysis of co-precipitation of fibrin(ogen) and Spike amyloid fibrils. Fibrinogen was covalently labeled with fluorescein and Spike amyloids were covalently labeled with Cy5. A, B, C) The fluorescence spectrum from a 25 µm^2^ region of interest (ROI) with a median spectral signature for each sample type. The top two panels depict the fluorescence spectra for each individual component. Fibrinogen and Spike amyloid fibrils were mixed, incubated for 200 minutes and were pelleted by centrifugation at 20 000 g (panel 3). A separate mix of Fibrinogen and Spike amyloid fibrils were subjected to thrombin cleavage and plasmin degradation and pelleted as above (panel 4). D) quantitative analysis of 27 ROIs collected from 3 different objects on the microscope slide. The ratio between fluorescence intensity at 516 nm and 688 nm describes the degree of co-precipitation of fluorescein labeled fibrin(ogen) and Cy5 labeled Spike amyloid fibrils respectively. Box indicates 25%-75% interval, whisker indicates mean ±1,5 stdev, central line indicates median and central open box indicates mean. Two-sample T-test was performed: ns= not significant; p>0,05=*; p>0,0001=****

Qualitative analysis of the experiments where fibrinogen was co-precipitated with Spike601 and Spike685 amyloid fibrils respectively, using confocal microscopy, demonstrates that fibrinogen co-precipitates with Spike601 amyloid fibrils (**Fig. 5A)**, but was barely visible when co-precipitated with Spike685 amyloid fibrils (**Fig. 5B**). On the other hand, when these reactions were subjected to thrombin to induce fibrin clots and centrifuged, Spike 685 amyloid fibrils were encapsulated within fibrin droplets while Spike601 amyloid fibrils were distributed more evenly within a fibrin mesh (**Fig. 5C-D**).

**Figure 5.**
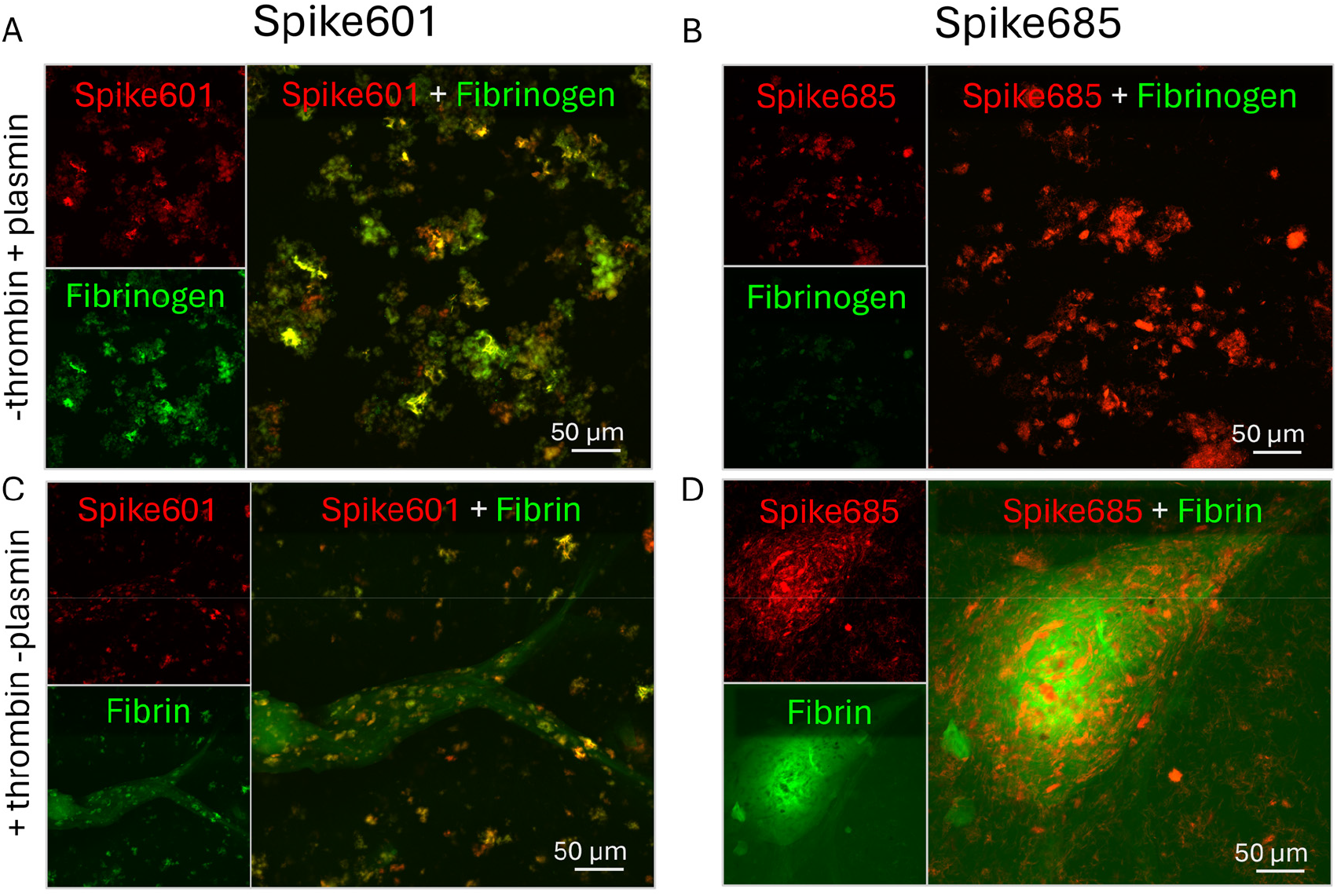
Representative confocal microscopy images for qualitative analysis of co-localization of fluorescein labeled fibrin(ogen) and Cy5 labeled Spike amyloid. Top panels illustrate samples where fibrinogen has been co-incubated with Spike amyloid prior to plasmin degradation. Lower panel illustrate the degree of co-localization between Spike amyloid fibrils and thrombin induced fibrin.

TEM analysis was performed to further investigate fibrinogen binding and fibrinolysis products in presence of Spike601 and Spike685 amyloid fibrils. Fibrinogen and Spike amyloids were co-incubated and sedimented by centrifugation. TEM analysis revealed that Spike601 amyloid fibrils in the presence of fibrinogen had a thin coating (**Fig 6B*i***) and also spheres of a size in accordance with the size of fibrinogen nodules (5-7 nm)[22] (**Fig 6B*ii***) attached to the fibrils. Co-incubation of fibrinogen and Spike685 amyloid fibrils on the other hand did not result in such coating. Spheres on Spike685 amyloid fibrils were scarcely found and on the contrary the apparent repulsion of the putative fibrinogen spheres from the perimeters of Spike685 amyloid fibrils was conspicuous (**Fig 6B*iii,iv***). Furthermore, *in vitro* formed and lysed clots were subjected to precipitation by centrifugation. In the absence of Spike amyloid fibrils, again spheres corresponding to the size of fibrin(ogen) nodules were visible (**Fig. 6C*i***, **D*ii***). If clots were formed in presence of Spike601, thin layers of amorphous material were deposited on the TEM grid (**Fig. 6D*ii***) and in it, slender amorphous aggregates were occasionally found (**Fig. 6C*ii,iii*, D*ii***). When clots were formed in presence of Spike685 amyloid fibrils, the pelleted material contained a large amount of dense granular structures with fibrous cores, sizing between ∼50 and ∼500 nm (**Fig. 6C*iv,v*)** and were evenly distributed over the TEM grid (**Fig. 6D*iii***). Spike amyloid fibrils were also found side by side to the granular structures (**Fig S2**).

**Figure 6.**
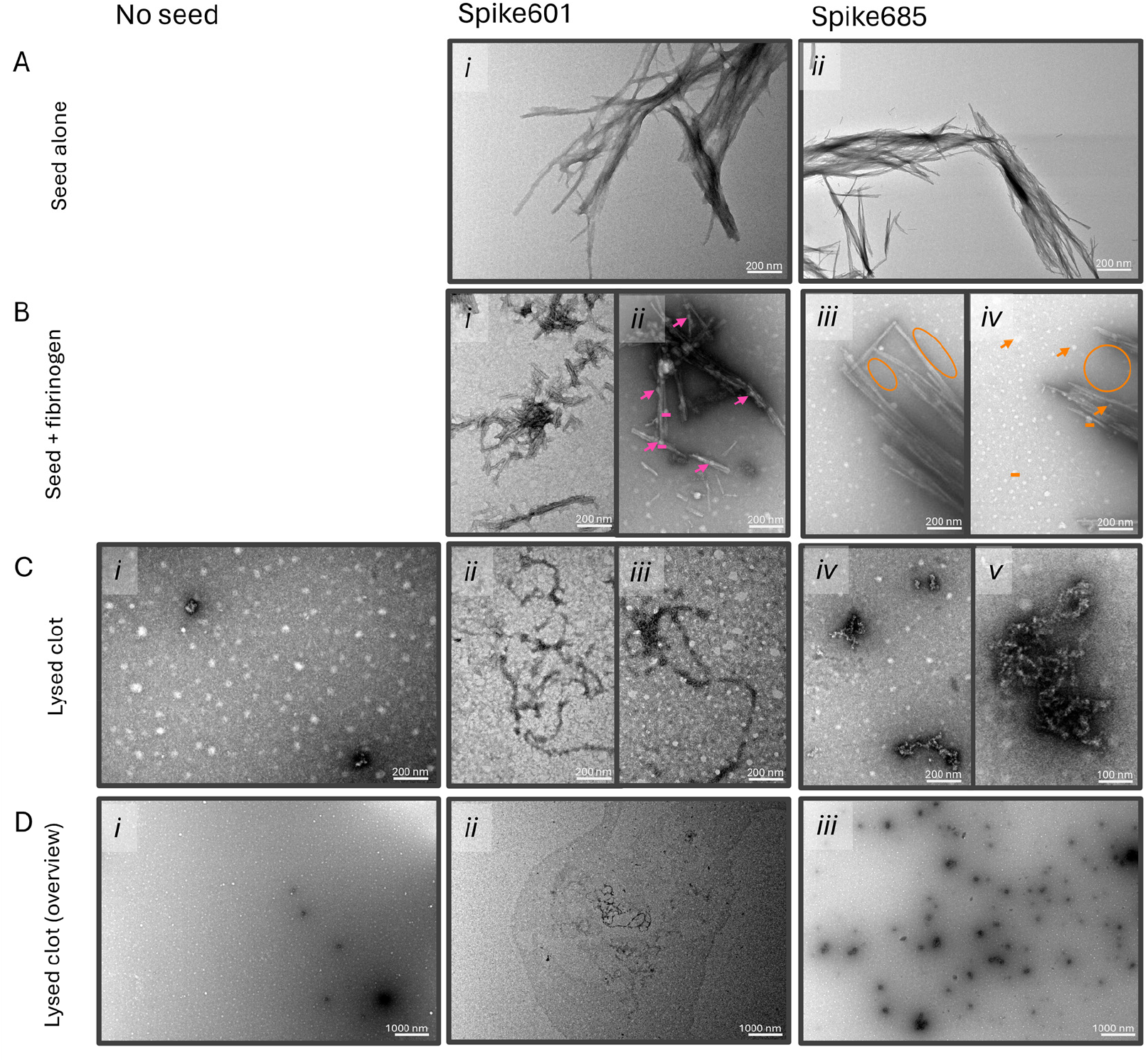
Representative negative stain TEM images showing A) Spike 601 and Spike685 amyloid fibrils B) Spike amyloid fibrils incubated with fibrinogen. Colored lines indicate the average diameter of the nodule parts of a fibrinogen molecule, arrows indicate suggested fibrinogen associated to amyloid fibril, encircled are areas depleted of suggested fibrinogen. C, D) Fibrin clots after lysis by plasminogen, in absence or presence of Spike amyloid fibrils. Spike amyloid fibrils visible side by side with fibrin colts exemplified in SFig 2.

## Discussion

COVID-19 is to a large extent a disease of the vasculature system and is associated with coagulation abnormalities in a large proportion of patients [23]. Insufficient fibrinolysis has been acknowledged as a prominent factor behind this phenomenon [24] and neutrophils have been pinpointed as a driving force of pathogenesis [25]. Events of thromboembolism has also been recorded as adverse event of COVID vaccination from several of the major vaccines on the market with the common factor being expression of SARS-CoV-2 Spike protein [26] [27]. Development regarding diagnostic markers of long COVID is also an area where research is focused [5]. A large number of potential biomarker candidates have been identified, but as of now no specific tests for molecular diagnosis of long COVID exists [28]. Furthermore, there are difficulties in distinguishing these potential biomarkers from other conditions. Microclots, suggested as a probable cause of neurological symptoms, have also been seen as a potential therapeutic target in long COVID [5] [29]. Fibrin degradation products such as D-dimer have been elevated in 25% of recovered COVID patients up to 4 months post infection [30].

In this work we focused on our previous findings of amylogenesis of fragmented Spike protein and its influence on fibrinolysis [10]. To this end we specifically studied if certain amyloid fibrils from different Spike sequences affected the fibrinogen-fibrin-fibrinolysis pathway.

### Fibrinogen, fibrin and fibrinolysis

Soluble fibrinogen is a haemostatic 340 kDa glycoprotein circulating in plasma at ∼2-5 mg/ml. Fibrinogen is enzymatically converted to fibrin upon vascular injury [31]. Structurally, fibrinogen is a dimer of three disulfide-linked polypeptide pairs: Aα, Bβ, and γ, forming a molecule with two D-domains and a central E-domain connected by coiled-coil regions [32] [33]. Fibrin network formation occurs in three stages [33]. First, thrombin cleaves the Aα and Bβ chains in the E-domain, releasing fibrinopeptides A and B and exposing “knobs” A and B, which bind to complementary “holes” a and b in the D-domains of adjacent molecules. Second, this interaction drives linear assembly into double-stranded protofibrils (∼500–600 nm). Finally, protofibrils laterally and longitudinally aggregate into thick fibrils and a 3D fiber network [31] [33]. Clot stability is enhanced by blood cell incorporation and covalent crosslinking via Factor XIIIa [34]. Fibrin clot structure is strongly influenced by the conditions during fibrin formation. A fibrin network can also be conformationally converted by imposed strain and partially convert from α-helical to β-sheet secondary structure which resulted in increased binding of the amyloid dye ThT and conversely a lower affinity for tissue plasminogen activator (tPA) [35].

Fibrinolysis is the regulated degradation of insoluble fibrin networks, crucial for prevention of thrombosis [36] [37]. The process is mediated by plasmin, a protease derived from its inactive zymogen, plasminogen. Plasminogen binds to lysine residues on fibrin fibers, where it is cleaved and activated by tPA. Activated plasmin then cleaves fibrin transversely, releasing D-dimers and other fibrin degradation products (FDPs) [36] [38]. The rate of fibrinolysis is determined by factors such as the fibrin fiber diameter, clot density, and pore size of the formed fibrin clot. A dense fibrin structure consisting of thinner fibres has been observed to lead to lysis resistance, as diffusion of tPA and plasminogen in the fibrin network becomes stalled.

### Fibrin amyloid microclots in long COVID

Fibrin amyloid microclots (fibrinaloids), 1-200 µm in size, have been suggested to be a main contributor to long COVID pathology [39]. The makeup of these fibrinaloids from *ex vivo* samples have been described to be primarily composed of fibrin with amyloid characteristics, but also other proteins such as α-2-antiplasmin, SARS-CoV-2 virion particles, and inflammatory molecules [8] [40]. Several recent reports illustrate that SARS-CoV-2 Spike protein influences coagulopathy. *In vitro* studies of blood plasma, using fluorescence microscopy to observe formed aggregates with amyloid dyes thioflavin T and Amytracker revealed amyloid components in Spike induced microclots *in vitro* [40]. The amyloid formation of these fibrinaloids *in vivo* have been argued to stem from interactions between fibrinogen and other proteins, such as the SARS-CoV-2 Spike protein or the acute phase protein serum amyloid A (SAA). Because of the β-sheet amyloid structures of fibrinaloids, and incorporation of proteolysis inhibitors (2)-anti-plasmin and plasminogen activator inhibitor 1, a resistance to normal fibrinolysis of fibrin clots is formed. This fibrinolytic resistance and aggregation of fibrin then leads to the formation of persistent microclots that suggestively cause long COVID symptoms. A consequence of fibrinaloids could be blockage of microcapillaries, which induces tissue ischemia and hypoxia. As fibrinaloids also incorporate inflammatory molecules and other proteins, it is suggested that these microclots could also contribute to the observed thrombotic endotheliosis in long COVID. Fibrin bound to the SARS-CoV-2 Spike protein generated proinflammatory blood clots that drive systemic thromboinflammation [41]. In addition, these proinflammatory fibrin clots are resistant to degradation and may possess a potential thromboembolic threat.

An association of microclots in general to cardiovascular disease (CVD) has also been made [42]. Changes in fibrin clot structures could be linked to different CVD risk factors, such as prothrombic disease phenotypes and hypertension. Up to 500 different clot-bound proteins were also determined to be included in CVD related plasma fibrin clots. Additionally, denser fibrin clot networks and hypofibrinolysis pose an increased risk of CVD [42].

The common components of SARS-CoV-2 infection and most regimens of COVID vaccination is the expression of Spike protein and the recruitment of neutrophils to the site of infection or injection. Brogna and colleagues used mass spectrometry utilizing the vaccine specific Proline mutation as bench mark and could establish expression of vaccine mediated Spike protein in 50% of tested samples and for up to 187 days post vaccination [43]. mRNA is detectable in blood up to 28 days after vaccination [44] and in the axillary lymph nodes and in myocardium up to 30 days in patients that died from heart failure in close temporal proximity to receiving mRNA vaccination against COVID [45]. A recent study used *in situ* hybridization to detect persistent vaccine mediated as well as SARS-CoV-2 virus induced Spike protein expression in the intima of cerebral arteries in over 40% of vaccinated patients who suffered haemorrhagic strokes [46]. Expression of Spike protein could persist up to 17 months after the latest vaccination as shown by immunohistochemistry. According to the same study, Spike protein positive histology was also found in unvaccinated cases and even without previous medical records of SARS-CoV-2 infection, indicative of mild or asymptomatic infection [46]. The concert of these studies is that Spike protein can be persistently expressed in several cell types and organs including cerebral arteries for an extended period after SARS-CoV-2 infection and mRNA vaccination generating Spike protein.

Amyloids in a more general context are known to interact with and impair the coagulation and fibrinolysis cascades [47]. SARS-CoV-2 Spike protein comprises several amyloidogenic sequences and we have demonstrated that SARS-CoV-2 Spike protein, when cleaved by neutrophil elastase forms amyloid and this is dictated by at least 7 amyloidogenic segments within the large Spike protein [10]. In addition, the Pretorius and Kell groups have demonstrated that addition of Spike protein to platelet poor plasma renders formation of microclots with amyloidogenic properties such as ThT positivity and fibrillar structure [39] [48].

With this background using pure *in vitro* systems, we herein scrutinized a rather complex molecular mechanism of how Spike derived amyloid fibrils could influence microclot formation. Our data suggest that specific Spike derived amyloid fibrils impacted fibrin folding and assembly and thereby cause impaired plasmin mediated fibrinolysis. Under the conditions used here (lower concentrations than in previous work [10] five out of seven Spike amyloid fibrils did not substantially affect fibrinogen-fibrin-fibrinolysis, whereas two affected the process in different ways. A closer analysis of the turbidity curves in the presence of Spike601 and Spike685 amyloid fibrils revealed that Spike601 amyloid fibrils delayed the formation of a fibrin clot but did not impair the fibrinolysis. Spike685 amyloid fibrils (and to some extent Spike365 amyloid fibrils) on the other hand enhanced the rate of fibrin clot formation and prevented the full removal of the formed fibrin clot by plasmin degradation. A titration of these amyloids revealed a decrease in fibrin formation rate as Spike601 amyloid fibrils concentration increases while a significant decrease in fibrinolysis was evident with addition of more Spike685 amyloid fibrils. Spike601 amyloid fibrils attracted fibrinogen, potentially blocking thrombin accessibility. Spike685 amyloid fibrils on the other hand had lower affinity for fibrinogen but stabilized fibrin formation and facilitated fibrin misfolding. Hence, fibrin formed in the presence of co-aggregating Spike685 amyloid fibrils resulted in hampered plasmin mediated fibrinolysis. It is notable that the amyloidogenic sequence encoding Spike685 is located directly after the furin cleavage site (at position 686) between the S1 and S2 regions of the prefusion state of the Spike protein. While this cleavage is not fully sufficient to dissociate S1′ and S2, this occurs after processing at S2′ (position 816) [49], it proposes a vulnerable dynamic nicked Spike protein state (**Fig. 7**). The 2P (K986P and V987P) mutations have been inserted in the vaccines to stabilize the prefusion conformation. However, it has been hypothesized that the same furin processing with shedding of S1 may occur after vaccination with the original Pfizer-BioNTech and Moderna mRNA constructs that contains the furin cleavage site [50]. Such a vulnerable aggregation protein state, as illustrated in **Fig. 7**, if presented on the cell surface would facilitate intermolecular interactions with neighbouring cleaved Spike proteins and Spike protein fragments from neutrophil elastase cleavage [10], facilitating misfolding, aggregation and recruitment of extracellular fibrin(ogen) proximal to cells expressing the Spike protein. This process may be an initiation seed for a persistent amyloidogenic microclot thrombus. It is important to note that many mutations have occurred in the viral SARS-CoV-2 spike protein since the original Wuhan strain, which we studied herein, several of which are in and around the furin cleavage site [51] and in other positions mainly in the RBD. This may have implications for the risk of microclot formation from SARS-CoV-2 infection. So far the main attributed cases of long COVID have been reported from the initial strains of the virus and possibly from some original vaccine constructs. Emerging innovative screening technologies enabling high throughput studies of putative patients with microclots was recently proposed such as nailfold capillaroscopy [52], which would be imperative for making clinical trials of therapeutic interventions.

**Figure 7.**
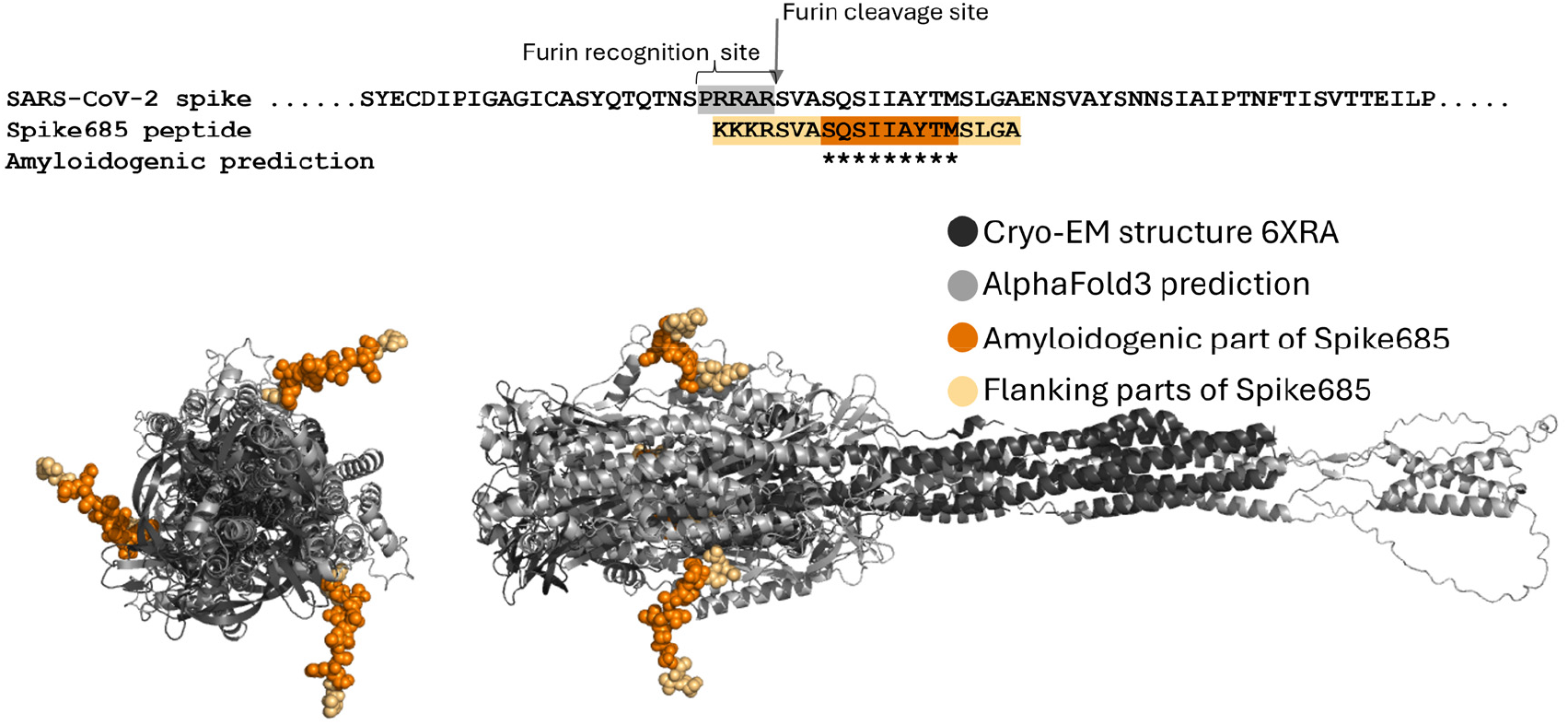
Hypothetical model of a vulnerable aggregation prone state of SARS-CoV-2 Spike protein. Sequence of the S1/S2 boundary with the highlighted furin cleavage site, compared with the Spike685 peptide (top). Structure model from AlphaFold3 (https://alphafoldserver.com/) of the full sequence of the S2 trimer overlayed with the cryo-EM structure of the post-fusion state of S2 (PDB:6XRA)[53], in top view and side view. The protein segment for Spike685 is highlighted in orange and protrudes out from the globular domains and appears accessible for intermolecular interactions.

In conclusion, our findings demonstrate how the amyloidogenic properties of SARS-CoV-2 Spike protein can have bearing on several of manifestations found in the complex multitude of symptoms of severe and long COVID, as well as of adverse side effects of Spike protein expression vaccination strategies. This study should foster further exploration of Spike protein targets such as Spike685 sequence containing Spike protein aggregates when mining for biomarkers of long COVID. The knowledge should also be considered for future development of vaccines to avoid exposure to amyloidogenic segments which may negatively influence human health.

## Materials and methods

### Spike amyloid formation

Amyloids of peptides derived from SARS-CoV-2 Spike protein (Wuhan strain) were generated as previously described [10]. In short, amyloidogenic amino acid stretches were detected using the online available WALTZ algorithm [19]. The peptides were custom synthesized by Genscript, NL, and delivered as lyophilized powder. The powder was resuspended in 100 % hexafluoro-isopropanol (HFIP) to a final concentration of 10 mg/ml. 1 mg/ml samples in PBS with 10% residual HFIP and 0.5 mg/ml samples with 5 % HFIP were subjected to 37 °C and shaking in glass vials for 24 hours to generate amyloid fibrils. Spike peptide sequences can be found in **Figure 1**.

### ThT spectra

Fibrils formed in glass vials were used for the fibrinogen experiments. To ensure fibril content, 10 µl aliquots in triplicate was added to 90 µl ThT in PBS to reach a final concentration of 0.1 mg/ml Spike peptide and 2 µM ThT in a 96-well black, untreated half area plates with flat transparent bottom (Corning costar 3880). Wells with ThT buffer supplemented with 10 µl fibrillation buffer (PBS+HFIP) was used as reference. ThT intensity was measured bottom up using a Tecan Infinite M1000 Pro plate reader, excitation wavelength 440 nm and emission scan every 5 nm between 470 and 600 nm.

### Congo red microscopy

Aliquots from the unstained fibrillation reactions were prepared for evaluation of Congo red birefringence as follows: 10 μl of 0.5 mg/ml Spike-peptide fibrils was added to 90 μl of Congo red staining solution (10 μM Congo red in 25 mM Tris-HCl, pH 7.5) resulting in a molar ratio of Spike peptide:dye 2.5:1. Stained fibrils were left to self-sediment over-night at 20 °C and were briefly centrifuged at 2000 rpm in a benchtop centrifuge. 3 μl from the bottom of the pelleted samples were transferred to superfrost gold glass slides (Thermo Fisher, Walldorf, Germany) and allowed to dry. The dried droplets were covered with fluorescence mounting medium (Dako, Glosrup, Denmark). Congo red stained samples were analyzed using a Nikon bright field microscope equipped with polarizers for both incoming light and in front of the detector.

### ThT kinetics

ThT measurement of amyloid formation over time (ThT kinetics) was performed as previously described [10] but at a peptide concentration of 0.5 mg/ml. In short, HFIP solubilized peptide was added to ice cold phosphate buffered saline (PBS) pH 7.4, supplemented with ThT to reach a final concentration of 0.5 mg/ml peptide, 2 μM ThT and 5% HFIP. Reference samples without peptide were run in parallel. The samples were distributed in 96-well black, untreated half area plates with flat transparent bottom (Corning costar 3880) placed on ice. The sealed plate was placed at 37 °C in a Tecan Infinite M1000 Pro plate reader with linear shaking between measurements with amplitude 2 mm and frequency 654 rpm. ThT intensity was monitored by bottom read mode with excitation at 440 nm and emission at 500 nm every 5 minutes for 24 hours.

### TEM of Spike amyloid fibrils

TEM grids from each of the Spike amyloid fibril samples were prepared from the fibrillation reactions as follows: 5 μL of samples were placed on 400 mesh carbon coated copper grids (Carbon-B, Ted Pella Inc.) and incubated for 2 minutes. Excessive salt was rinsed of by one wash with 5 μL milliQ water. The grids were negatively stained with 2% uranyl acetate for 30s before blotted dry and air dried overnight. Transmission electron microscopy (TEM) imaging was performed using a Jeol JEM1400 Flash TEM microscope operating at 80kV.

### Turbidity monitoring of fibrin formation and fibrinolysis

A turbidity based fibrin formation-fibrinolysis experiment was performed inspired by Terasawa, Okumura and colleagues [54]. In detail:

Frozen fibrinogen stock solution (−20 °C) of 25 mg/ml in dH_2_O (human, Merck, 341578) was thawed at 37 °C in a heating block. A Mastermix of 0.625 mg/ml fibrinogen and clot buffer (20 mM 2-[4-(2-Hydroxyethyl)piperazin-1-yl]ethane-1-sulfonic acid (HEPES) pH 7.4, 0.12 M NaCl, 1 mM CaCl_2_) was prepared and kept at 37 °C. Thrombin working-solution (WS) 1 U/ml in clot buffer was prepared using a refrigerated thrombin stock solution of 500 U/ml (bovine, Merck, 605157-1KU) and kept cold. A lysis mixture of 1 µM plasminogen, 2.4 nM tPA in clot buffer was prepared using a stock solution of 50 µM plasminogen (human, Roche, 10874477001) and 1,2 nM tPA (Human, Merck, T0831-100 μg) and was kept on ice during fibrin clot formation.

Each turbidity experiment was performed in six replicates. Because of the short deadtime of the experimental set up, each plate run contained two Spike seeded experiments (eg Spike192 and Spike258 amyloid fibrils) together with one unseeded control used as reference for experiments on that plate.

Thrombin-induced fibrin clot formation was performed in a Corning Costar 3881 plate (non-binding, half-area well with total maximum well volume of 190 µl). First, 60 µl of Mastermix (0.625 mg/ml fibrinogen) was applied to each assigned well in a pre-warmed microplate (37 °C). 1.5 µl of Spike amyloid fibrils (1 mg/ml) was added and mixed in the corresponding wells of the two Spike variants. The plate was then inserted into a Fluostar Galaxy microplate reader (BMG Labtech) and the experimental run was started, first measuring baseline turbidity at 355 nm. After 4 cycles of 45 seconds (3 minutes) the measurement was paused, and the plate was taken out.

To each well containing Mastermix, 15 µl of thrombin WS (1 U/ml) was applied and mixed in with pipette, discarding the pipette tip for every well. The final clot concentrations of fibrinogen and thrombin were 0.5 mg/ml and 0.2 U/ml, respectively. After addition of thrombin WS the plate was reinserted to the plate reader, and turbidity measurement at 355 nm resumed for 60 minutes at 37 °C. After formation of fibrin clots, 75 µl of lysis mixture consisting of 1 µM plasminogen and 2.4 nM tPA was applied in each well by carefully dispensing the lysis solution against top of plate walls. The final concentrations of plasminogen and tPA were 0.5 µM and 1.2 nM, respectively and the final Spike amyloid fibril concentrations were 10 µg/ml. Turbidity at 355 nm was then measured at 37 °C for 2.5 hours (150 minutes) in a final well volume of 150 µl.

### Dose dependent fibrin formation and fibrinolysis

A dose dependence experiment was performed for the most interesting Spike amyloid fibrils: Spike601 and Spike685. Here increasing concentrations of Spike amyloid fibrils were added, to reach final concentrations 1.4, 2.8, 5.6, 10, and 20 µg/ml of Spike amyloid fibril per experiment. Each fibril concentration was run in 4 replicates, including also 4 replicates of unseeded control.

First, 60 µl of Mastermix (0.625 mg/ml fibrinogen), thrombin WS (1 U/ml thrombin), and lysis mix (1 µM plasminogen, 2.4 nm tPA) were prepared as previously described for the turbidity assay using the Fluostar Galaxy microplate reader (BMG Labtech). 0.1 mg/ml Spike fibril solution was prepared by mixing 10 µl of 1 mg/ml Spike amyloid fibrils to 90 µl of PBS, and vortexed. Starting with the lowest concentration of 1.4 µg/ml, 2.1 µl of 0.1 mg/ml Spike fibrils was added and mixed to each corresponding well. Similarly, for Spike fibril concentrations 2.8 µg/ml and 5.6 µg/ml, 4.2 µl and 8.4 µl of 0.1 mg/ml Spike amyloid fibril solution were applied to assigned wells. Lastly, for Spike amyloid fibril concentrations 10 µg/ml and 20 µg/ml, 1.5 µl and 3 µl of 1 mg/ml Spike amyloid fibril solution was mixed into corresponding wells. After measuring baseline turbidity, fibrin clotting was initiated as previously described adding 15 µl of thrombin WS to each well. The final concentrations of fibrinogen and thrombin were 0.5 mg/ml and 0.2 U/ml, respectively. Turbidity measurement of clot formation was then run for 60 minutes. After completed fibrin clot formation, fibrinolysis was started by applying 75 µl of lysis mix to all wells, and the decrease in turbidity was observed for 150 minutes. The final concentrations of lysis mix were 0.5 µM plasminogen 2.4 nm tPA.

### Fibrin clot formation for fluorescence microscopy

Cy5 labeling of preformed Spike amyloid fibrils of Spike365, KKK-Spike601, and Spike685 (made as described above) was performed by first dissolving 1.5 mg of Cyanine5 NHS ester (Cy5) (Lumiprobe, Westminster, MD, USA) in 100 µl DMSO. This stock was further diluted 10-fold and 10 µl was added to 500 µl Spike fibrils 0.5 mg/ml dissolved in PBS buffer pH 7.5 with 5% HFIP. This resulted in a final concentration of 45 µM Cy5 and 227 µM Spike peptide. Staining was allowed to proceed for 1 h, whereafter the reaction was quenched with 10 µl 0.1 M Tris-HCl pH 8. The samples were diluted with 1000 µl PBS and were concentrated and diluted in two additional rounds of buffer change to PBS buffer in Amicon Ultra Centrifugal Filters, 10 kDa MWCO (Millipore). Cy5 labeled KKK-Spike amyloid fibrils were stable for about one week in room temperature.

Fluorescein labeled fibrinogen was generated by dissolving Fluorescein-5-Isothiocyanate (FITC) (F1906, Invitrogen) in DMSO and mixing with 1 ml of 0.625 mg/ml fibrinogen in clot buffer (20 mM 2-[4-(2-Hydroxyethyl)piperazin-1-yl]ethane-1-sulfonic acid (HEPES) pH 7.4, 0.12 M NaCl, 1 mM CaCl_2_) kept at 37 °C. This resulted in a final concentration of 1 µM FITC and 1.8 µM fibrinogen for the labeling reaction. Labeling was allowed to proceed for 1 h. One part FITC-labeled fibrinogen was mixed with 10-fold excess unlabeled fibrinogen for use in the fibrin and fibrinolysis experiments as described in that section in the presence of Cy5 labeled Spike amyloid fibrils.

For the fluorescence microscopy experiments, 60 µl per well of freshly FITC-labeled fibrinogen diluted with unlabeled fibrinogen (total 0.625mg/ml) was mixed with 3 µl of the three different Cy5 labeled Spike amyloid fibrils and were subjected to thrombin cleavage and lysis by plasminogen/tPA in a plate reader at 37 °C as described in fibrinogen-fibrin-fibrinolysis experiments. Samples that were not thrombinated were kept in parallel in the plate reader to establish the affinity between fibrinogen and the Spike amyloid fibrils. Samples before addition of thrombin and after thrombin cleavage and plasmin lysis were collected and sedimented by centrifugation 10 minutes, 20 000 g at 10 °C. The pelleted material was subjected to hyperspectral imaging with an epifluorescence microscope (Leica 6000B equipped with a spectral cube, ASI, Migdal Ha Emek, Israel) and with fluorescence confocal microscopy (Zeiss 780 LSM-system) [21].

## Acknowledgements

We thank the ProLinC core facility and the core facility at the medical faculty of Linköping University for instrumentation access.

This study was funded by grants from Olle Enqvists Stiftelse, Swedish Research Council (2023-03931) and Gustav V and Drottning Viktorias Foundation.

## Supporting information

**Figure S1.**
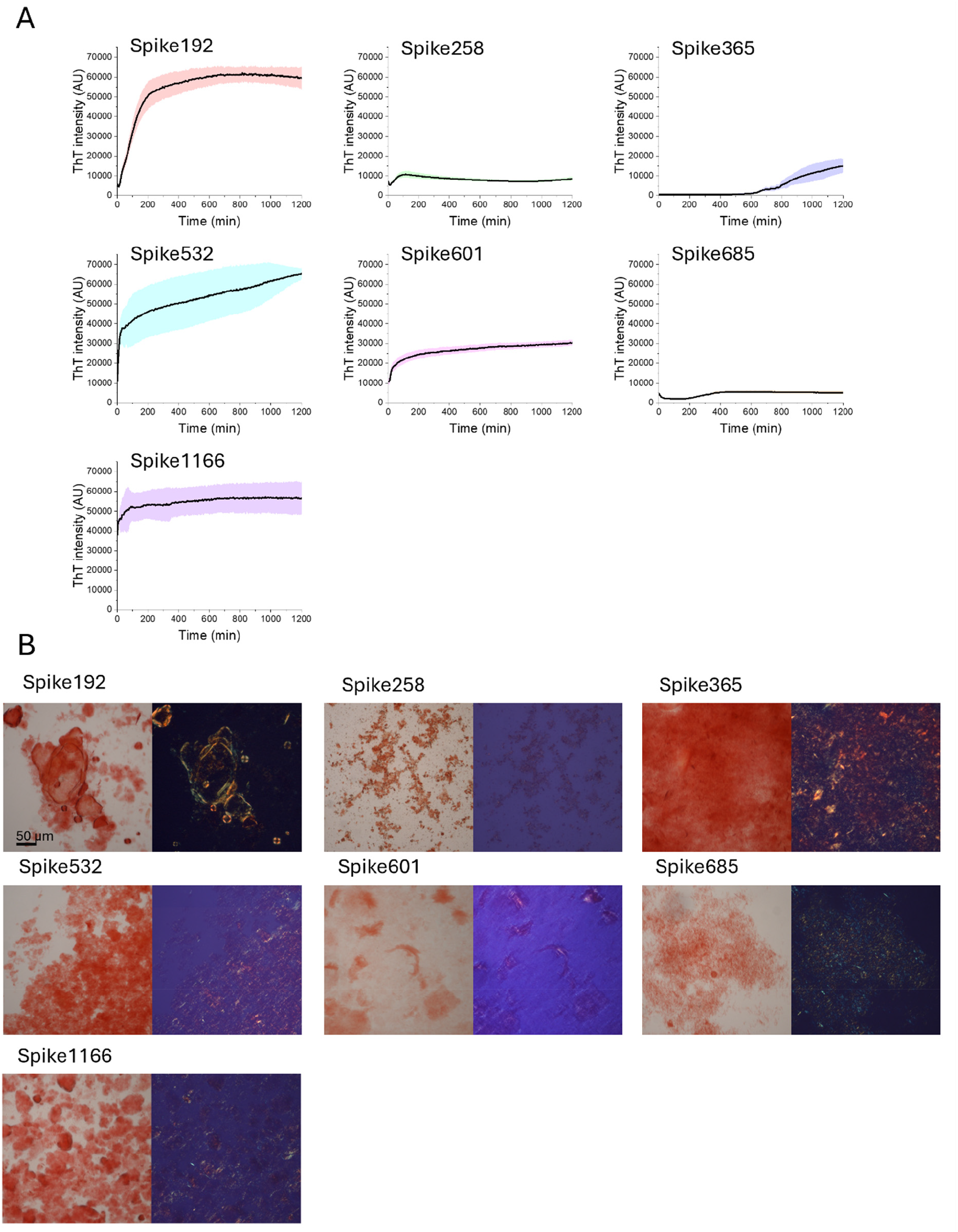
A) Amyloid conversion of the seven Spike peptides was monitored by ThT intensity increase over time. B) Amyloid content at the endpoint of the Spike peptide fibrillation reaction was determined by light microscopy of Congo red stained samples without (left panel) and with (right panel) crossed polarizers. Fibrils were generated with spike peptide concentration of 0.5 mg/ml, in 5% HFIP in PBS buffer pH 7.5, 37 °C.

**Figure S2.**
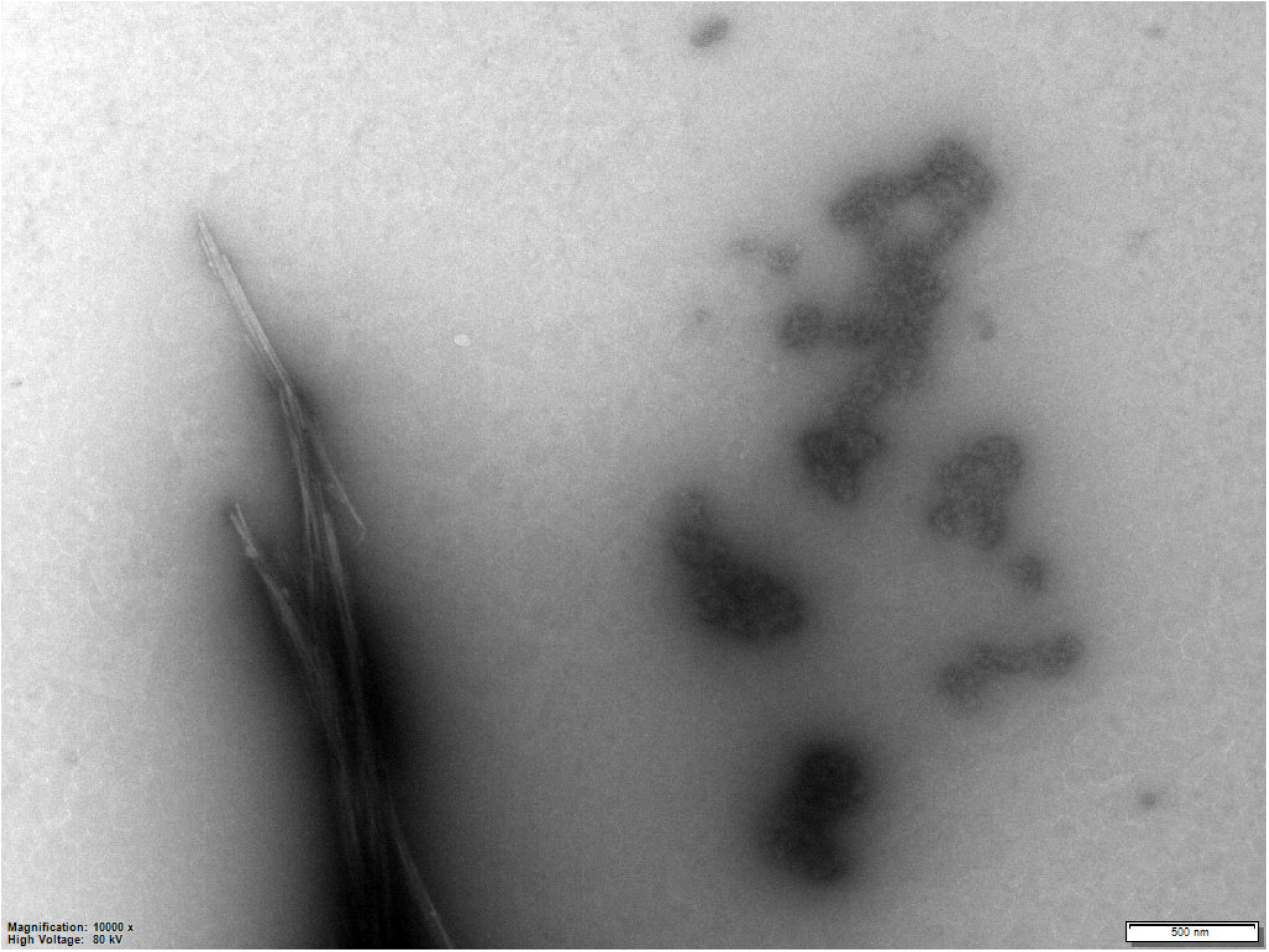
Spike amyloid fibrils were visible side by side with granular fibrin clots on the TEM grids of samples with lysed clots exemplified here by Spike685 amyloid fibrils to the left and granular fibrin clots to the right.

